# MtDNA d-loop variation in Prespa dwarf cattle (*Bos taurus*) from Albania

**DOI:** 10.1101/2023.03.30.534932

**Authors:** Angela Schlumbaum, Jose Granado, Gruenenfelder Hans-Peter, Joerg Schibler

## Abstract

Albanian Prespa dwarf cattle belong to the Busha group of cattle with short horns, uniform coat color and low withers heights. Their breeding population is very small and the population is recognized as endangered. Here we report mtDNA d-loop diversity (475bp) from 10 unrelated Albanian Prespa individuals from Liqenas, one male and nine females. Diversity measures are high and members of the taurine haplogroups T3, common in Europe, and one female with T1, common in Africa and rare on the Balkans, were detected. This supports a complex female genetic history in the past.

## 1. Introduction

Cattle from the Balkans are often geographically isolated and frequently belong to endangered breeds or strains. Some populations do not even have a breed status, and it was suggested to use the term “strains of a metapopulation” instead [1]. This is the case with Albanian Prespa dwarf cattle which belong to the so-called Busha group (Buša), comprising short-horned (brachycerous), small cattle with withers heights between 95cm and 115cm [2] and which are classified as at risk by DAD-IS (http://www.fao.org/dad-is/browse-by-country-and-species/en/). This type of cattle corresponds e.g. to small cattle during the Neolithic Horgen culture, the Iron Age and mediaeval period in the alpine foreland [3] [4] [5] and may as thus give a hint about pre-historic cattle populations. Albanian Prespa cattle have a history of isolation with few events of admixture recently [6] [2]. Analyses based on mtDNA d-loop sequences, STRs and whole genome SNP data [7] [8] [9] [2] [1] support the special status of the Busha cattle and recommend their conservation. To our knowledge, no mtDNA d-loop data from Albanian Prespa are publicly available, and therefore composition and diversity of female lineages remain unclear.

## 2. Materials and Methods

Tissue samples from 10 unrelated Prespa cattle were collected from local owners by H.-P. G. in Liqenas, Albania in 2008 (Tab. S1) in the framework of SAVE (Safeguard for agricultural varieties in Europe) activities. DNA was extracted with Qiagen Blood and Tissue Kit (Qiagen), PCR was carried out with primer targeting pos. 16018 – 207 of the reference sequence V00654 (BRS) (Anderson et al. 1982) and directly sequenced by Microsynth, Balgach, CH [10] (for details see supplemental text). Sequences are registered with EMBL with accession numbers: MW080668 – MW080677.

## 3. Results and Discussion

Based on 16 polymorphic positions (SNPs) 8 different haplotypes were found. Only one haplotype, T3a, was shared between individuals 3, 15, and 35. Other 5 individuals belong to different variants of haplogroup T3, of these two are variants of the BRS haplotype T3b (Table S1, Figure 1).

**Figure 1.**
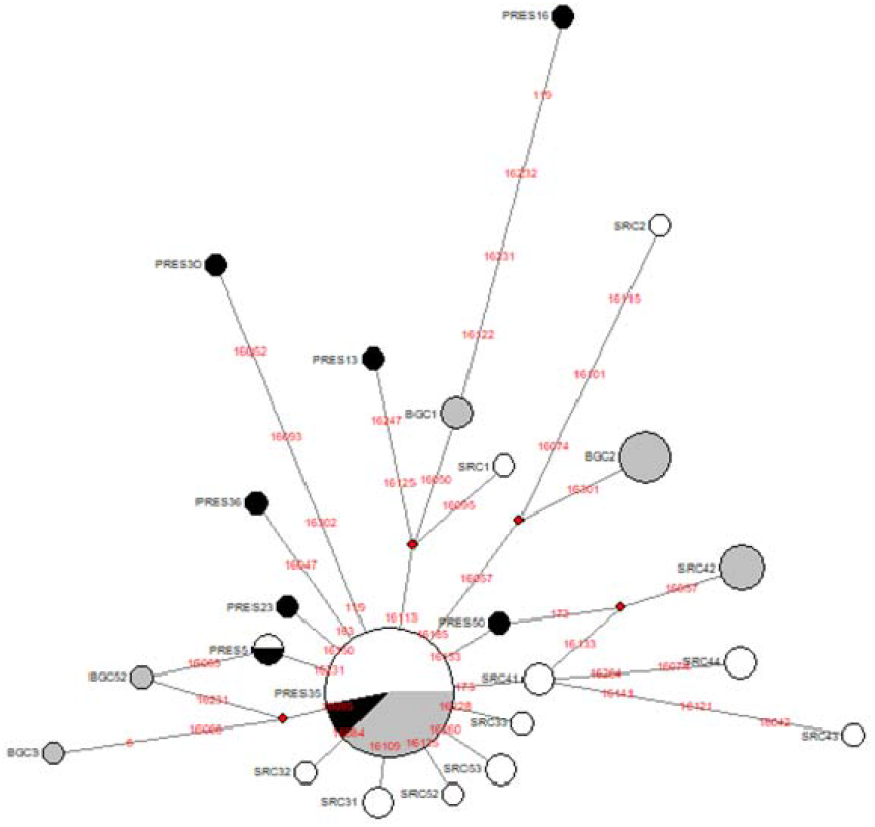
Median-joining network of Prespa cattle (black dots) in comparison with Bulgarian Grey (grey dots) and Shorthorn Rhodopean (white dots) (data from [13]). Pos. 16255 and 169 are e cluded.

Two cattle affiliate with haplogroup T1. In particular, Prespa16 belongs to T1 based on the diagnostic SNPs at positions g.16050C>T, g.16113T>C, g.16122A>G and g.169A>G. Prespa13 may also belong to T1, missing the diagnostic mutation at pos. g.16050C>T [11]. Within the network (Fig. 1) both individuals are located on the branch leading to T1 haplotypes, however Prespa13 falls within the T3 haplogroup variation, if the dataset is expanded (e.g. from [12], data not shown). Note that none of the two sequences carries pos. g.16255T>C, considered to be a backward mutation [11]. Given that haplotype Tb3 is the most frequent taurine haplotype in Europe, our data confirm that cattle were indeed not maternally related and represent herd diversity. African mtDNA T1 haplotypes in Prespa cattle were not detected in earlier surveys of Busha cattle [7] [13] but are present in low frequencies in other cattle populations in the Balkans, Greece, Italy or Anatolia [14] [13]. This indicates contacts between these regions at any time in the past or that this individual is a rare relic/descendant from the initial spread of cattle which reached the region about 6100 cal BC, e.g. [15]. We did not detect neither Q, nor T2 haplotypes (Fig. 1) or T6 haplotypes. The network shows little haplotype sharing between Prespa and the two other Balkan breeds. The main T3 haplotype is shared between all 3 breeds, in addition Prespa5 shares one haplotype with SRC [13].

Haplotype diversity and MNPD (mean number of pairwise differences) of Prespa cattle are high and comparable to the SRC population (Tab. 1). The high number of private haplotypes is reflected in negative Tajima’s D (Tab. 1) and might indicate a recent bottleneck. However, *P* is not significant on a 5% level. The *F*_ST_ value of 0.144 (*P*=0.0) supports structuring between the 3 populations, probably due to isolation and different population histories.

**Table 1.** Diversity measures of Prespa cattle. * data from Hristov et al. 2015 Diversity measures of Prespa cattle. * data from Hristov et al. 2015 Diversity measures of Prespa cattle. * data from Hristov et al. 2015

| | N | hts | htdiv | MNPD ($\pi$) | Tajima's D | P |
| --- | --- | --- | --- | --- | --- | --- |
| Prespa | 10 | 8 | 0.93 ± 0.07 | 3.82 ± 2.09 | -1.5 | 0.06 |
| Shorthorn Rhodopean* | 19 | 12 | 0.94 ± 0.03 | 4.07 ± 2.12 | -1.11 | 0.13 |
| Bulgarian Grey* | 40 | 7 | 0.73 ± 0.04 | 1.53 ± 0.93 | -1.03 | 0.14 |

Altogether, the results of this study show high genetic diversity derived from partial mtDNA d-loop sequences along with a presumably complex female history of the Prespa cattle breed. Our data may aid in conserving female diversity in endangered Prespa cattle strains with a small breeding population [8] [6]..

## Statement of conflict of interest

We declare no conflict of interests

## Author Contributions

Conceptualization, A.S. and J.S.; Collection and lab work, J.G. H.-P.G; Writing – Original Draft Preparation, A.S., J.G., J.S., H.-P.G.

## Notes

### Competing Interest Statement

The authors have declared no competing interest.

